# Beyond the Triple Helix: Exploration of the Hierarchical Assembly Space of Collagen-like Peptides

**DOI:** 10.1101/2024.05.14.594194

**Authors:** Le Tracy Yu, Mark A. B. Kreutzberger, Maria C. Hancu, Thi H. Bui, Adam C. Farsheed, Edward H. Egelman, Jeffrey D. Hartgerink

## Abstract

The *de novo* design of self-assembling peptides has garnered significant attention in scientific research. While alpha-helical assemblies have been extensively studied, exploration of polyproline type II (PPII) helices, such as those found in collagen, remains relatively limited. In this study, we focused on understanding the sequence-structure relationship in hierarchical assemblies of collagen-like peptides, using defense collagen SP-A as a model. By dissecting the sequence derived from SP-A and synthesizing short collagen-like peptides, we successfully constructed a discrete bundle of hollow triple helices. Mutation studies pinpointed amino acid sequences, including hydrophobic and charged residues that are critical for oligomer formation. These insights guided the *de novo* design of collagen-like peptides, resulting in the formation of diverse quaternary structures, including discrete and heterogenous bundled oligomers, 2D nanosheets, and pH-responsive nanoribbons. Our study represents a significant advancement in the understanding and harnessing of collagen higher-order assemblies beyond the triple helix.

## Introduction

The *de novo* design of self-assembling peptide systems is of great interest to scientists. Among the various assemblies described to date, structures assembled from α-helices are the most extensively studied resulting in precise design capabilities through rational and computational approaches.^1–13^ In addition to α-helices, researchers have explored the self-assembly of β sheets^14–17^, hybrid assemblies^18,19^, and the 3_10_ helix^20^. The Polyproline type II (PPII) helix is a protein secondary structure frequently found in collagen, the most abundant extracellular protein.^21,22^ However, compared to other secondary structures, PPII helices have been underexplored. In collagen, PPII helices are characterized by the Xaa-Yaa-Gly sequence repeat, with three PPII helices associating into a collagen triple helix.^23^ In this super helix, the three peptide strands are offset from one another by 3n+1 amino acids, where n is an integer. This offset defines a leading, middle and trailing peptide strand and is termed the helix’s “register”.^24^ Several structural studies of collagen triple helices have been conducted to develop synthetic collagen-mimetic materials with the polymeric higher-order assemblies being the most well-developed as reported by the Conticello group^25–29^, the Raines group^30^, the Xu group^31^ and our group^32^. However, achieving precise control over the self-assembled state to the degree which has been achieved in α-helical coiled coils remains elusive due to limited understanding of the sequence-structure relationship in collagen higher-order assembly. For instance, computational algorithms like Alphafold2 and Rosetta, while highly effective in predicting α-helical assemblies, encounter difficulties in accurately predicting collagen oligomer formation.^6,33–35^

Previously we introduced an approach to creating a hollow bundled collagen oligomer assembly using peptides derived from the immune protein component C1q.^36^ This family of proteins, which also includes SP-A, ficolins, and manose binding lectins, is characterized by a bouquet-like structure, with the stem region consisting of bundled collagen triple helices, a branching region and a globular region. ^37^ Specifically, the C1q stem forms a hexameric assembly of triple helices, with each subunit being a heterotrimer containing three different peptides: A, B, and C. Through the synthesis and characterization of short peptides derived from the stem region of C1q we explored the self-assembly mechanism of this region. We found that the formation of the hollow (ABC)_6_ octadecameric bundle is independent of the C-terminus collagenous and globular fragments as well as disulfide bonding near the N-terminus. However, a short N-terminal non-collagenous domain is necessary for effective assembly of the (ABC)_6_ complex.^36^ This work demonstrated the structure of the first discrete (non-polymeric, monodisperse) oligomeric collagen triple helical assembly, providing a valuable model for future investigation of inter-triple helical interactions. While some studies have suggested particular registrations of C1q’s triple helical domains^38,39^, challenges associated with the characterization and ambiguities introduced by discontinuities in the Gly-Xaa-Yaa repeat leave the register of the three peptides uncertain. This poses a challenge to the investigation of amino acid sequences critical for the formation of this quaternary structure. In this work we turn our attention to another protein within the C1q family, Pulmonary Surfactant Protein A (SP-A). In contrast to C1q, SP-A features a homotrimeric collagen-like stem region, consisting of only one type of peptide. This characteristic dramatically simplifies the study of the sequence-structure relationship in hierarchical assemblies of collagen-like peptides, facilitating structural studies.

We dissected the sequence derived from the stem region of SP-A protein and synthesized these relative short, collagen-like peptides. Upon self-assembly, we have successfully constructed a hollow octadecameric bundle similar to that observed for C1q. Subsequently, we conducted mutation studies to locate amino acid sequences critical for oligomer formation. Amino acid sequences including hydrophobic Isoleucine residues and the charged Arginine and Aspartic acids, were found to favor the oligomer formation. Furthermore, we identified the N-terminal noncollagenous domain as critical for protein assembly, as peptide mutants lacking this domain failed to form discrete collagen oligomers. The sequence-structure relationship information was applied to guide the de novo design of collagen-like peptides to form bundled structures of triple helices. We used a (Gly-Pro-Hyp)_10_ peptide as a template starting point in the design, which has been known to create a stable collagen triple helix, but without higher order oligomerization.^40–42^

Amino acid residues that facilitate interhelical charge-pair interactions and hydrophobic packing were introduced into the design peptides. We have successfully obtained hierarchical assemblies of four different architectures including discrete oligomeric bundles, heterogeneous oligomeric bundles, polymeric ribbons and 2D nanosheets, showcasing the potential for forming diverse quaternary structures with collagen-like peptides (Fig. 1). This study demonstrates peptide sequences conforming to a collagen-like sequence provide a rich space for hierarchical assembly, providing new approaches for future synthetic design as well as insight into natural collagen hierarchical assembly.

**Fig. 1.**
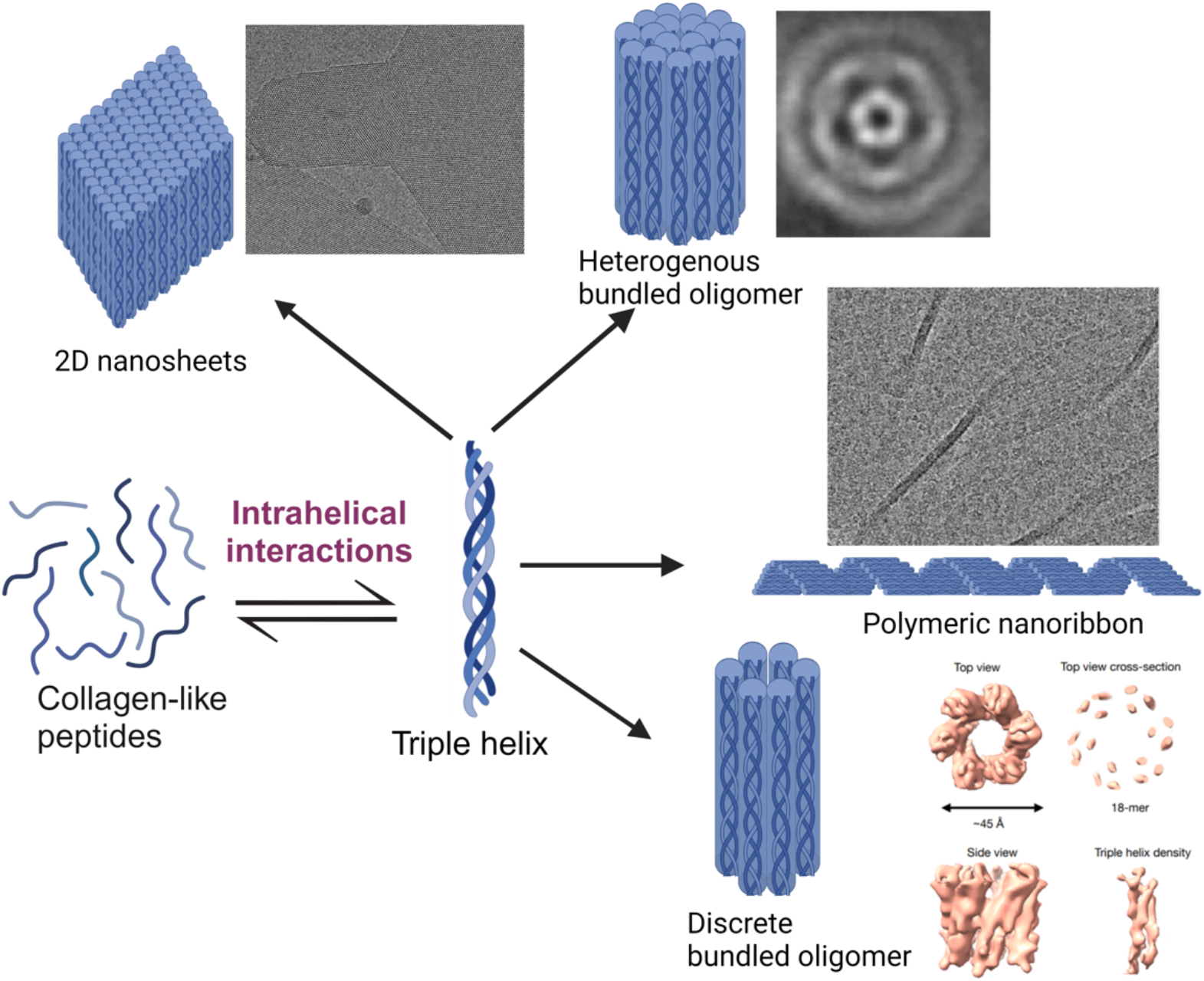
Schematic illustration of the diverse hierarchical assemblies formed in this study using collagen-like peptides.

### Collagen-like Homotrimer and Octadecamer Formation

We started with amino acid sequence analysis of protein SP-A. We obtained the full-length amino acid sequence of natural SP-A and highlighted the collagen-like region by identifying the glycine residue in every third amino acid in the sequence, and recognized the discontinuity of the (Gly-Xaa-Yaa) sequence pattern starting from Gly45 (numbering starts after the signal sequence).^43^ The N-terminal 1-44 amino acid sequences ahead of the discontinuity are therefore assumed to be the stem collagen-like region (see Fig. S1 for sequence dissection details).^43^ Similar to C1q, disulfide bonded dimer formation has previously been observed at the Cys6 position in protein SP-A.^44^ Our previous study of C1q protein suggests that the higher-order assembly is independent of the disulfide bond. We therefore designed peptides Cys6Ala mutants of protein SP-A for self-assembly study.

We conducted an analysis of the amino acid sequences of SP-A protein from various organisms, revealing differences in the amino acid sequences of the N-terminal noncollagenous domain. To investigate the structural influence of the noncollagenous region, we made three collagen-like peptides for self-assembly studies. The first peptide, SPA-bcb, contains the collagen-like region exclusively (Fig. 2a). The other two peptides, SPA-Human and SPA-Rat are derived from the stem region of the human SP-A α2 chain and the rat SP-A protein, respectively (Fig. 2a). All these peptides were synthesized using solid-phase peptide synthesis, purified via reverse-phase HPLC, and characterized using mass spectrometry (refer to Supplementary Information Table S1 and Fig. S2-S13 for details).

**Fig. 2.**
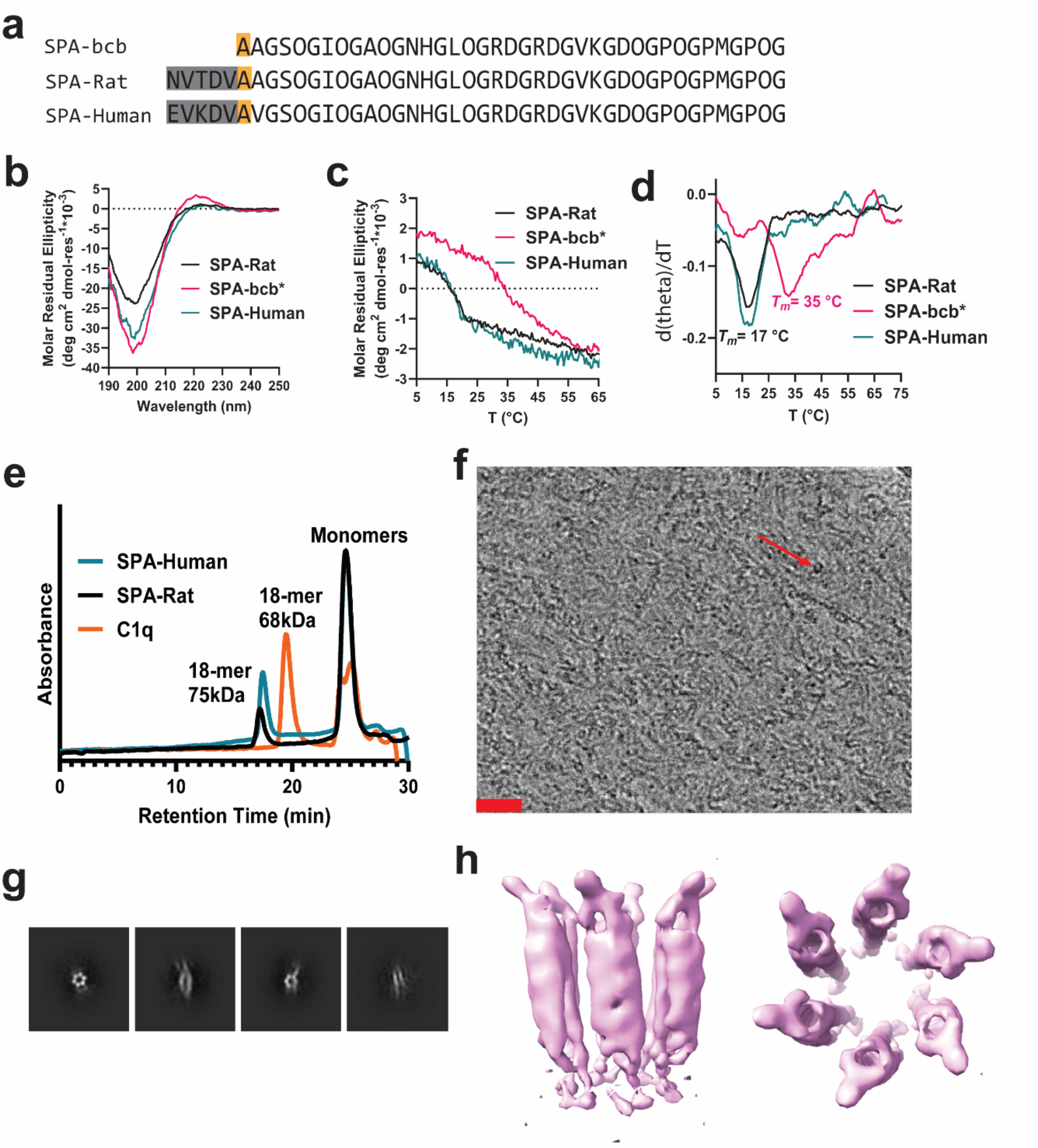
Characterization of SP-A derived peptide assemblies. a) Amino acid sequences of SP-A derived collagen-like peptides SPA-bcb, SPA-Rat, and SPA-Human. Grey highlighted the non-collagen-like domain, and yellow highlighted the alanine residue, which is cysteine in the native pulmonary surfactant protein-A. Yellow highlights Cys6 to Ala mutations. Numbering starts after the signal sequence. b) CD spectra of the folded triple helices. c) Thermal stability study of the assemblies. Signals were monitored at 224 nm. *SPA-bcb peptides formed a heterogeneous assembly that contains precipitate. d) The first-order derivatives of the melting curves presented in panel c). e) Size exclusion chromatography of SPA-Rat, SPA-Human and C1q 18-mer. f) Representative cryo-EM image of SPA-Human peptide assembly. The scale bar is 20 nm. The red arrow highlights the hollow protein nanoparticles. g) 2D class averaging processing of the cryo-EM images of SPA-Human peptide assembly. h) Low-resolution density map of SPA-Human peptide oligomeric assembly.

After folding under buffered conditions, the peptide assemblies were characterized for their secondary structure using circular dichroism (CD) and oligomeric state using size exclusion chromatography (SEC) and cryo-EM (Fig. 2). Peptides SPA-bcb, SPA-Rat and SPA-Human all present a characteristic collagen triple helix feature in CD spectra showing a positive signal at approximately 224 nm (Fig. 2b). The thermal stability of the triple helices assembled by peptide SPA-Rat and SPA-Human are comparable, having a melting temperature (*T*_*m*_) of 17 °C, which indicates that the sequence difference near the N-terminus of these two proteins did not alter the triple helix folding stability (Fig. 2c, d). The SPA-bcb peptide precipitated upon folding and remained insoluble in different solvent systems such as aqueous buffers, isopropanol, or 2,2,2-Trifluoroethanol. Such turbidity impacted CD measurements and resulted in an increased signal-to-noise level. According to the CD melting experiments, the pepetide aggregation derived from SPA-bcb has a *T*_*m*_ of 35 °C. The sample solution became clear above this indicated melting temperature. We hypothesize that the aggregation may enhance the thermal stability of individual triple helices. While we suspect that this precipitation may consist of polydisperse triple helical aggregates, Cryo-EM characterization of the SPA-bcb sample only showed amorphous features (Fig. S14).

Size exclusion chromatography (SEC) was used to characterize the oligomeric state of the soluble peptide assembles of SPA-Rat and SPA-Human. The octadecameric assembly previously derived from C1q peptides was used as a point of comparison^36^ in addition to a collagen triple helix standard^45^ and globular protein standards (Fig. S16). SEC of both human and rat SP-A peptides revealed peaks eluting at 25 minutes corresponding to monomeric peptides and at 17 minutes corresponding to an octadecameric assembly with a molecular weight near 75 kDa (Fig. 2e). The SEC results confirmed that SPA-Rat and SPA-Human peptides associate into homogenous collagen-like oligomers. The equilibrium of the 18-mer formation is favored in the SPA-Human compared to sample SPA-Rat, indicating that the difference in the N-terminal amino acid sequence of these two peptides affects oligomer formation (Fig. 2a). These results demonstrated that the oligomer formation in SP-A is independent of the C-terminal region of the protein and the N-terminal disulfide bonds . The presence of the short noncollagenous domain, however, prevents protein aggregation and favors the discrete oligomer assembly. These are similar to results we had observed for C1q.^36^

We further characterized the octadecameric peptide assembly of SPA-Rat (Fig. S15) and SPA-Human using cryo-EM (Fig. 2f and S17). The raw cryo-EM images of SPA-Human peptide assembly show hollow protein particles of ∼ 6 nm in diameter (Fig. 2f) as well as more filamentous assemblies (Sup. Fig. S17). The 2D class averages show side views and top views of the hollow oligomeric peptide assemblies (Fig. 2g). Similar cryo-EM results were observed for SPA-Rat (Fig. S15). The size of the particles in the SP-A human sample appeared rather heterogenous, especially when looking at the side views. This could be because some of the assemblies’ side views were actually filaments containing multiple particles. The overall heterogeneity complicated three-dimensional structure determination. Despite this, a low-resolution electron density map was obtained after 3D reconstruction with C6 symmetry imposed (Fig. 2h). We observed features similar to the previously published structure of the C1q peptide assembly. Given the similarities to the C1q structure and the low resolution (∼6 Å) triple helical features, we believe that this structure is representative of a portion of the SP-A peptide assembly.

### Mutation Studies Reveal Critical Sequence-Structure Relationship

The supramolecular octadecameric assembly from the SP-A stem region provided a model for studying the interhelical interactions that drive the collagen triple helices to oligomerize. We next conducted point-mutation studies to investigate the amino acids critical for higher order assembly. We focused our preliminary screening on the charged and hydrophobic amino acid residues. To investigate the effects of charge-pair interactions on the oligomer assembly, we made peptide mutants of Arg24 in the Xaa position to be proline and Asp25 in the Yaa position to be hydroxyproline residue since the (Gly-Pro-Hyp)_n_ sequence is regarded as the canonical sequence for collagen assembly (Fig. 3a). The samples were prepared for self-assembly and then subject to CD and SEC characterizations. Fig. 3b,c and S18a show the CD characterization of the SP-A mutant peptides. A homotrimeric helix formed with a *T*_*m*_of 19 °C for sample SPA-D-O but no triple helix formation for sample SPA-R-P was observed. Despite the fact that proline is known to be more stabilizing for triple helices than arginine^46,47^, the Arg24 to proline mutation eliminated self-assembly, indicating that Arg24 is critical in some way for the folding of the octadecameric assembly. SEC results of SPA-R-P peptide only show peaks of monomer species while SPA-D-O peptide assembly suggests formation of trimer and oligomer (Fig. 3d). The oligomeric peak in sample SPA-D-O assembly is broad and shifted to a lower mass region compared to the 18-mer peak, indicating that the assembly decreased in size. This suggests that aspartate also plays an important role in helix oligomerization, possibly through inter-triple helical interactions.

**Fig. 3.**
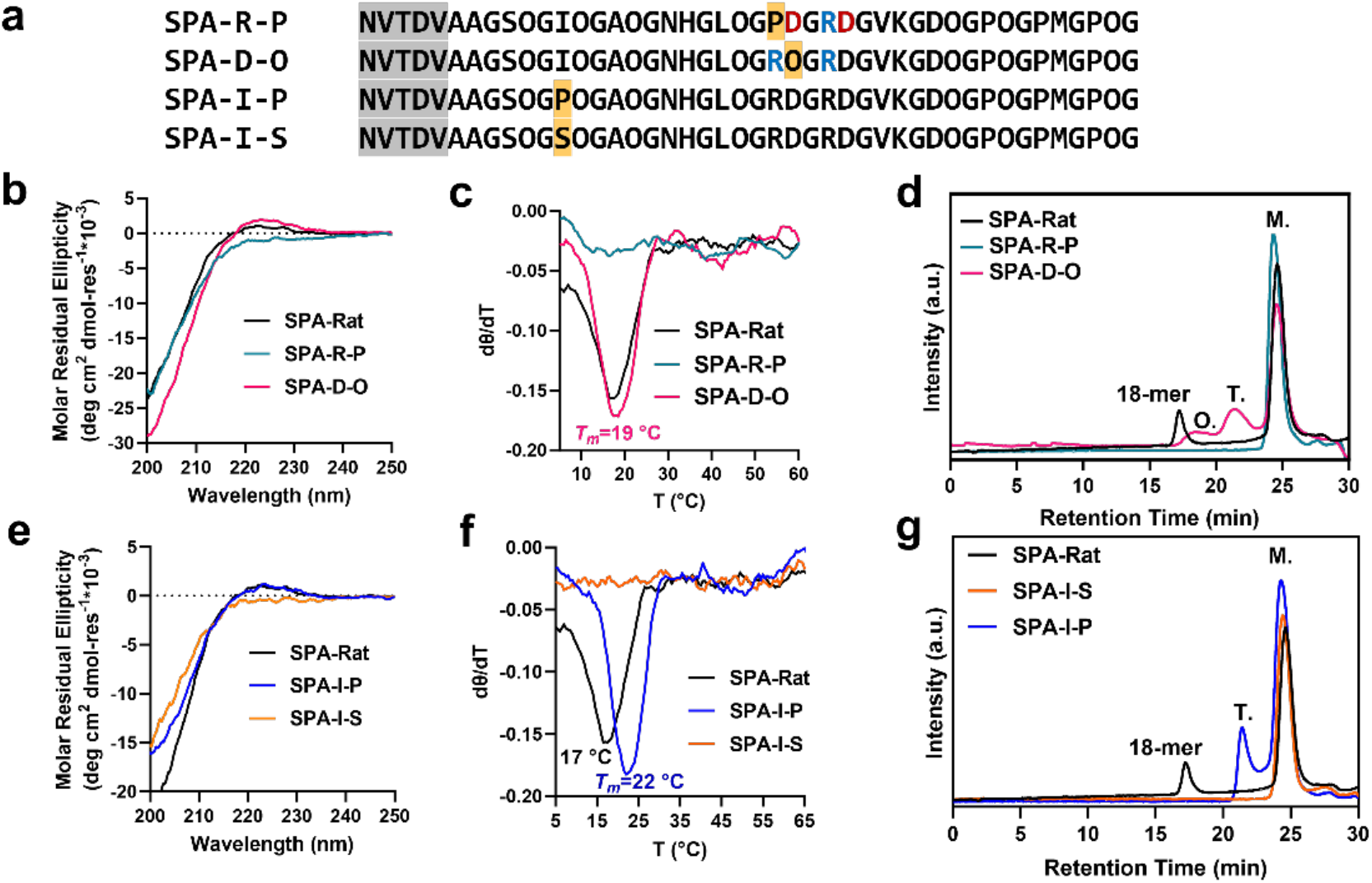
Mutation study of SP-A derived peptides. a) Amino acid sequences of the mutant collagen-like peptides derived from SPA-Rat peptide. b) CD spectra of peptide mutants involving Arg24Pro and Asp25Hyp mutations. The amino acid sequence numbering starts after the cleavage of the signal peptide.^43^ c) Thermal stability study of the mutant peptides SPA-R-P and SPA-D-O. d) SEC results of peptide mutants SPA-R-P and SPA-D-O. e) CD spectra of the SPA peptides involving Ile12Pro and Ile12Ser mutations. f) Thermal stability study of the mutant peptides SPA-I-P and SPA-I-S. g) SEC results of peptide mutants SPA-I-P and SPA-I-S. “M.” represents monomer and “T.” represents trimer.

We additionally investigated hydrophobic residues since hydrophobic packing is one of the main interactions driving folding and self-assembly as illustrated by α-helical coiled-coils. To test the importance of hydrophobic packing in collagen oligomerization, we replaced the conserved isoleucine (Ile12) residue with the canonical proline (SPA-I-P) or hydrophilic serine (SPA-I-S). The CD spectra of both are presented in Fig. 3e, indicating triple helix formation in peptide SPA-I-P, but not I-S. The CD melts further confirmed our assumption. No thermal transition was observed for peptide SPA-I-S sample in the CD melting plot while a *T*_*m*_of 22 °C was observed by peptide SPA-I-P (Fig. 3f and S18b). SEC characterization of sample SPA-I-P suggests triple helix formation but no higher-order oligomerization (Fig. 3g). Even though the Ile to Pro mutant has enhanced thermal stability and favored triple helix formation, it did not promote oligomerization. This suggests that oligomerization is not correlated with triple helix formation, a critical finding. Replacement of this hydrophobic residue with a hydrophilic Serine ablated triple helix assembly completely: the CD spectrum lacked the PP2 signature, and only monomeric species were observed in the SEC (Fig. 3e, f, g). The point-mutation studies together revealed that charged-pair interactions involving arginine and aspartate and hydrophobic interactions may exist to favor collagen oligomerization.

### Design of pH-responsive Nanoribbon Formation

Directed by the sequence-structure relationship information revealed in the mutation studies, we prepared collagen-like peptides designed to assemble beyond the level of a triple helix guided by the insights gained from SP-A. The (GPO)_10_ peptide sequence was used as a template for the design (Fig. 4a).^41,42^ We added a short noncollagenous domain of “DAA” at the N-terminus of the peptide as the presence of noncollagenous domain reduces unwanted aggregation and favors a more specific oligomer formation. The choice of a charged residue “Asp” in this region decreased the pI point of the peptide to approximately 4, lower than the physiological pH that peptides were prepared to fold. This was anticipated to have a similar effect as the natural non-collagenous sequences which also have a net negative charge. In the SP-A peptides, the hydrophobic residues Ile, Leu, Val and Met were spaced out by three triplets. We therefore introduced the same types of hydrophobic amino acid residues into the design peptide in the *i, i+9, i+18* and *i+27* positions (Fig. 4a, DES-H peptide). “RDGRDG” charged sequences were introduced into the middle of the peptide for electrostatic interactions, similar to the SP-A peptides (Fig. 4a, DES-1 peptide). These peptides were synthesized and characterized in the same fashion as previous peptides. The CD spectra of the peptides show a characteristic signal of 225 nm, indicative of triple helix formation (Fig. 4b). The thermal stability of the formed triple helices was found to be 37 °C, and 49 °C for the DES-1 and DES-H triple helices, respectively (Fig. 4c). The folded DES-1 peptide assembled into hollow protein nanoparticles in MES buffer, pH 6.1 as demonstrated in cryo-EM images (Fig. S22). Further data processing suggests that these hollow particles are a mixture of three different assemblies: a relatively rare doublet ring structure in addition to much more common bundles of five and six collagen triple helices (Fig. 4d-g). The resolution information of the structure is listed in Table S2. This indicates that the predicted sequence-structure relationship investigated in the mutation study correctly identified key elements controlling higher order assembly. Whether the doublet structure was made up of hexamers, pentamers, or a combination of each was impossible to determine at the low resolution obtained. No higher-order assembly was observed with the DES-H peptide, indicating that the hydrophobic residues are insufficient to drive assembly alone and that the charged residues included in DES-1 are critical.

**Fig. 4.**
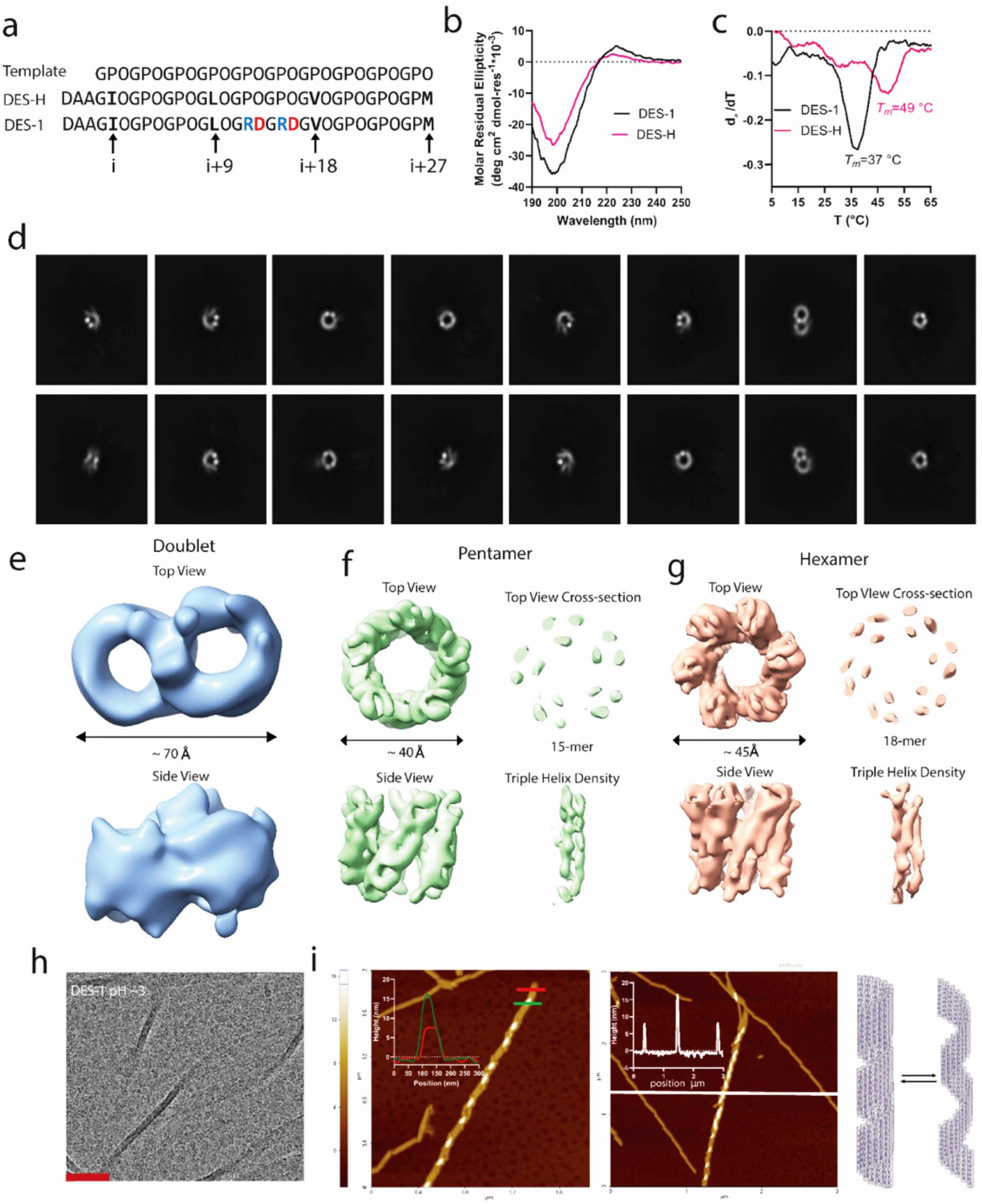
Structural characterization of the pH-responsive DES-1 peptide assembly. a) Amino acid sequence of template peptide (GPO)_10_ and the design peptides. b) CD spectrum of the design peptide (left panel). c)The first-order derivative curve of the CD melting curves of the designed peptides. (right panel) d)) 2D class averages of the particles showing a range of classes including pentameric assemblies, assemblies with a doublet, and hexameric assemblies. The peptide assembles in pH 6.1. e) Low resolution partial cryo-EM structure of the doublet DES-1 particle assembly.f) Partial cryo-EM structure of the pentameric DES-1 assembly. The top view and side view of the structure are gaussian low pass filtered using a standard deviation of 1.5. The cross section and triple helix density are not filtered. g) Partial cryo-EM structure of the hexameric DES-1 assembly. The top and side views are low pass filtered using a standard deviation of 1.8. The cross section and triple helix density are unfiltered. h) cryo-EM image of the design peptide assembly at pH 3.5. DES-1 peptide has assembled into nanoribbon structure. The scale bar is 50 nm. i) AFM characterization of peptide assembly at pH 3.5. The cartoon representation of the equilibrium of the tightly and loosely packed nanotubes is displayed on the right.

To test our hypothesis that charged residues are critical for the protein oligomer assembly, we tested DES1 at different pH. We made a sample at pH 3.5, lower than the pKa of Asp, a sample at pH 6.1 which is the default pH we made our samples, and a sample at the mildly basic pH of

8.0. After 3 weeks of equilibration, the sample prepared at pH 3.5 became turbid. CD spectra in Fig. S19a suggests peptide DES-1 folded into collagen triple helices at all tested pH values. CD melting results displayed in Fig. S19b and c indicated that the melting temperatures of DES-1 peptide folded in pH 3.5, pH 6.1 and pH 8.0 are 31, 38 and 37 °C. The *T*_*m*_differences indicates that the assemblies in pH 6.1 and 8.0 is likely similar to each other but different from the assembly in pH 3.5. Cryo-EM characterization demonstrated that DES-1 peptide had undergone further assembly into nanoribbons with a width of approximately 50 nm and varied length at pH 3.5 (Fig. 4h and Fig. S20). The 2D class average of the cryo-EM results of the nanotube region in this DES-1 peptide assembly at pH 3.5 show a uniform diameter of 50 Å with a space between striations of 10 Å (Fig. S20c). Scanning AFM of this peptide at pH 3.5 exhibited similar nanoribbons (Fig. 4i and Fig. S21). The nano-ribbon assembly is heterogenous with two general architectures observed: tightly packed nanotubes and more loosely packed nanoribbons, indicating the possibility of equilibrium between the two assemblies. The nanoribbon structure has a height of approximately 9 nm and the double-layer region has a height of approximately 18 nm (Fig. 4i). The nanoribbon formation has slow kinetics and nanostructure assembly was observed only after at least 2 weeks of equilibration at room temperature (Fig. S23). These supramolecular polymers help to illustrate the role of the DAA non-collagenous domain and control over the isoelectric point of the assembly as the low pH employed here allows for neutralization of long range ion-ion repulsion and subsequent extensive assembly. The cryo-EM image of the sample equilibrated at pH 8.0 showed similar hollow protein particles as seen in the peptide assembled at pH 6.1 (Fig. S22). This is reasonable as there is no expected net charge difference of the peptide at pH 6.1 vs. pH 8.0. These results suggest that the DES-1 peptide assembly is pH-responsive and the aspartate side chain is involved in the hierarchical assembly process.

### *De novo* Design of Heterogeneous Oligomeric Bundle and 2D Nanosheets

The pH-responsiveness of the DES-1 peptide indicates that charged residues play a significant role in driving the higher-order assembly. Motivated by this information, we designed collagen-like peptides containing only charged amino acid residues, but with no other deviations from the standard (Gly-Pro-Hyp) repeat (Fig. 5a). RD1 and RD2 peptides contain a short non-collagenous domain to prevent unwanted aggregation and each peptide contains four triplets of the “RDG” sequence. The “RDG” sequence in the RD1 peptide is centered in the amino acid sequences whereas that of the RD2 is spaced out by four triplets of “GPO” sequences. Both peptides assembled into collagen triple helices as indicated by the CD spectrum signal of 225 nm as seen in Fig. 5b. However, the thermal stability of the two peptide assemblies varied significantly. The CD melting results of RD1 and RD2 show *T*_*m*_ values of 50 °C and 34 °C (Fig. 5c, d), respectively. It has been suggested that the thermal stability of collagen triple helices is related with the amino acid propensity.^23,46,48^ RD1 and RD2 have identical amino acids compositions and therefore might be expected to have similar thermal stability. By contrast, the melting temperatures vary significantly, suggesting that different higher-order assemblies are likely formed by these two peptides. Additionally, the melting curve of RD1 peptide has broad thermal transition, suggesting the higher-order assemblies formed by this peptide is heterogenous.

**Fig. 5.**
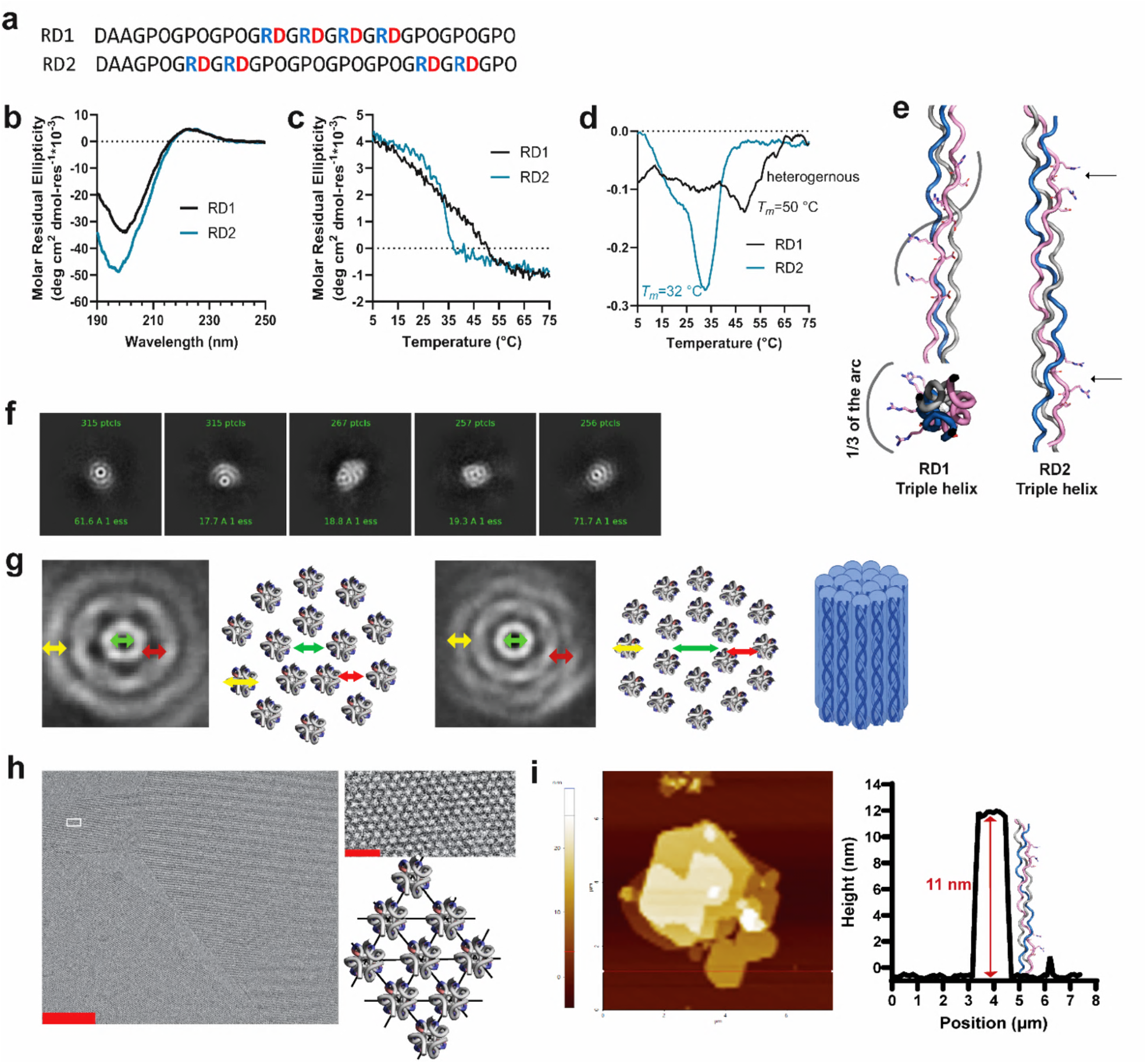
Characterization of RDG designed peptides. a) Amino acid sequences of designed peptides containing different “RDG” sequence pattern. b) CD spectra of design peptides. c) CD melting curves of design peptides. The signals were monitored at 225 nm. d) The first-order derivative curves of the melting curves displayed in c). e) Cartoon illustration of RD1 and RD2 triple helices. The model is created by amino acid substitution of PDB file 3b0s using pymol. The charged side chains occupy one third of the arc of the RD1 triple helix and the side chains of charged residues nearly align along the axis in the RD2 triple helix. For clarity, only charged residues of the leading chain are shown. f) 2D class averages of the RD1 peptide assembly from cryo-EM data. g) Measurements of the shelled ring structure and schematic illustration of the assembly. Yellow symbol measures the width of the ring (10 Å). Green symbol measures the size of the hollow pore (14 Å in the pentameric ring and 17 Å in the hexameric ring). Red symbol measures the distance between two rings (10 Å) h) cryo-EM characterization of RD2 nanosheets peptide assembly. The scale bar is 50 nm. The white square region was zoomed in and displayed on the right panel. The scale bar is 5 nm. i) AFM characterization of RD2 nanosheets peptide assembly.

Dynamic light scattering (DLS) of RD1 showed the formation of nanoparticles of approximately 19 nm (Fig. S24). Cryo-EM also demonstrated nanoparticles formation (Fig. S25). The 2D class averages of the cryo-EM data show hollow protein particles with multiple layers and a concentric pore (Fig. 5f). Most of the particles observed form either pentameric or hexameric structures in the innermost layer (Fig. 5g). The innermost layer of the hexameric structures has a hollow pore size of 17 Å, similar to the C6 assembly formed by C1q peptides.^36^ The width of each layer of the ring is approximately 10 Å, matching the diameter of a single triple helix (Fig. 5g). Based on this we assume that triple helices oligomerized into hollow pentameric or hexameric assemblies via charge-pair interactions between the side chain of arginine and aspartate in the RD1 peptides. Subsequently more triple helices bundled onto the oligomer, forming the second layer of triple helices. The unsatisfied charge-pair interactions likely initiate the formation of multiple layers of triple helices.

Different from RD1 peptide, the RD2 peptide formed two-dimensional nanosheets with varied size as observed from the cryo-EM images (Fig. 5h and S26). The width of each assembly unit is approximately 10 Å, corresponding to the width of a triple helix and the space between the striation of the nanosheets is approximately 14 Å, which may correspond to the distance between individual triple helices. AFM measurements of the 2D nanosheets suggest that the height of the nanosheets is approximately 10 nm (Fig. 5i and S27), which corresponds to the height of a triple helix. We hypothesize that the triple helices pack parallelly into 2D nanosheets via charge-pair interactions. Similar collagen triple helical 2D nanosheets, which like these lack patterened hydrophobic amino acids, have been reported by the Conticello group.^26,28,29^

In our investigation of SP-A peptides, we have highlighted the significance of the short noncollagenous domain. To elucidate this point, we specifically incorporated the “DAA” sequence into our peptide design. Our aim was to ascertain whether the presence of the “DAA” noncollagenous region truly influences the assembly process. To test this hypothesis, we developed a new peptide variant, RD2-no DAA (Table S1). By subjecting it to the same folding conditions as the RD2 peptide, we observed a notable difference in assembly behavior. The RD2-no DAA peptide formed 2D microcrystals exceeding 3 micrometers in size (refer to Fig. S28 and 29 for TEM and AFM), significantly larger than those formed by the RD2 peptide. This observation lends additional credence to our hypothesis, suggesting that the short noncollagenous domain may serve to deter aggregation or the formation of excessively large assemblies.

## Discussion

In this study, we have investigated three peptides derived from protein SP-A. Folding studies suggested the short non-collagenous region prevents aggregation that happened in SPA-bcb peptide. This precipitation/aggregation might also be caused by the fact that the peptides were assembled at a pH close to the isoelectric point (pI) as decreased solubility of proteins was observed when pH is close to the pI.^49^ However, we assume that the nocollagenous domain has additional underexplored structural functions that affect the bundled collagen-like oligomer folding. For example, we have observed that the oligomer formation has an enhanced equilibrium fraction in SPA-Human which differs from SPA-Rat only in the noncollagenous domain. The amino acids in SPA-Rat and SPA-Human at the 7^th^ position are alanine and valine, respectively. Valine is more hydrophobic than alanine, and we suspect that additional hydrophobic interaction may exist in the assembly of SPA-Human peptide to enhance oligomer formation.

From the characterization of the SPA-mutants, we observed no triple helix folding of SPA-R-P peptide but SPA-D-O mutant still form trimer and an oligomer of a different oligomeric status. This suggests that the Arginine side chain may form H-bonding with the carbonyl oxygen of the neighboring triple helix where the Asp residue was replaced. Therefore, oligomer formation was partially maintained in SPA-D-O but not SPA-R-P. Peptide mutant SPA-I-P gained enhanced thermal stability of the triple helix, but the oligomer formation was ablated, indicating that a stable triple helix will likely be more reluctant to form higher-order oligomers. We, therefore, do not think the formation of a triple helix is the prerequisite for oligomerization, rather, we believe that the oligomer formation happens simultaneously with the trimer formation. The fact that we did not observe trimer intermediate in SEC supported this hypothesis.

Interestingly, the DES-1 oligomers at pH 6.1 formed discrete structural populations of pentamers, hexamers, and doublets. The SPA-human cryo-EM structure also revealed hexameric structures in addition to heterogeneity. The discrete nature of these structures is likely a combination of the presence of both charged and hydrophobic interactions. The DES-1 structure with its two “back-to-back” repeating RDG motifs assembled into these discrete structural states while the addition of more RDG residues into the peptide sequence in RD1 resulted in the loss of discrete oligomeric states as additional “layers” of triple helical interactions appeared to form. Because our cryo-EM reconstructions were not of the entire complexes, as well as due to the limited resolution, we were unable to build atomic models of these assemblies. This limits our interpretation of the structural results. Despite this, it seems reasonable that the addition of more arginine and aspartate residues increases the charged-charged interaction between triple helices allowing for ordered layers outside of a hexameric, pentameric, or doublet assembly.

Serendipitously, the DES-1 peptide is a pH-responsive assembly. Previously, a pH responsive collagen assembly was developed by the Conticello group, using charged amino acid side chains of arginine and glutamate residues.^27^ The findings suggesting that charged residues can be included for future development of pH-sensitive materials.

RD1 and RD2 have formed different quaternary structures. Even though we do not have atomic level characterization of these assemblies, we believe that the difference between the sequence patterns is expected to result in the different three-dimensional geometry of the side chains in the triple helices. We have modeled a single triple helix of RD1 and RD2 peptides in pymol. The side chains of RDG sequences in RD1 peptide spiral up the helical axis from C to N term and occupy approximately 1/3 of the helical arc (Fig. 5e) versus the side chains of the RDG residues are on the same side of the triple helix (Fig. 5e). For clarity, only charged residues of the leading chain is shown. We hypothesize that in the folding process, the leading chain of one triple helix communicates with the leading chain of the neighboring triple helix, similarly for middle and lagging chains. Consequently, the differences in the tertiary structure of a triple helix will result in that the RD1 triple helices pack parallelly but arced whereas the RD2 triple helices pack parallelly and almost linear, forming hollow heterogeneous bundled oligomers and 2D nanosheets, respectively.

## Conclusion

In this study, we obtained discrete octadecameric assembly using collagen-like peptides derived from protein SP-A. Our exploration into the sequence-structure relationship of this oligomer formation involved a detailed examination of a variety of peptide mutants, particularly emphasizing the role of charged and hydrophobic residues in protein folding. We discovered that the charged residues of arginine and aspartate side chains, along with hydrophobic residue isoleucine, are key factors of facilitating oligomer assembly. Using this information, we designed collagen-like peptides that formed a pH-responsive collagen bundle-nanoribbon assemblies, heterogenous bundled oligomers and 2D nanosheets. This finding highlights the significance of amino acid sequence patterns in determining the 3D structure of the assembly, combined with previously published research on collagen-like nanofibers, underscore the versatility and complexity of collagen-like peptide assembly.^25,30–32^While our study does not fully exhaust the space of assembly possibilities, we anticipate that our findings will offer valuable insights for future efforts in protein design, especially concerning collagen-like mimetic peptides.

## Author Contributions

L. T. Yu: conceptualization, data curation, formal analysis, methodology, investigation, writing – Original draft. M. A. B. Kreutzberger: data curation, formal analysis, methodology, investigation, writing – review & editing. M. C. Hancu: data curation, investigation, writing – review & editing. T.H. Bui: methodology, writing – review & editing. A.C. Farsheed: methodology, writing – review & editing. E. H. Egelman: supervision, funding acquisition, writing – review & editing. J. D. Hartgerink: supervision, funding acquisition, writing – review & editing.

**Acknowledgement**

This work was funded in part by the National Science Foundation (CHE 2203937) and the Robert A. Welch Foundation grant (C-2141). EHE was funded by the NIH (GM122510). L.T. Yu would like to thank Dr. Wenhua Guo and Dr. Xu Wang for their help and discussion with sample characterization.

## Experimental Details

### Peptide Synthesis

All the reagents were obtained from Sigma Aldrich. Peptides were synthesized manually using solid-phase peptide synthesis with FMOC chemistry. In detail, a low-loading rink-amide resin was used to generate C-terminal amidated peptides. The resin was swelled twice with dichloromethane (DCM) and twice with N,N-dimethyl formaldehyde (DMF). 25 % v.v. piperidine in DMF was added into the resin to deprotect the FMOC protecting group for 5 mins. The resin was washed five times with DMF and tested with chloranil test or Kaser ninhydrin test to confirm presence of free amines. The amino acid, activating reagent hexafluorophosphate azabenzotriazole tetramethyl uronium (HATU) and a weak base diisopropylethylamine (DiEA) were dissolved in DMF in a molar ration of 1:4:4:6 (resin/amino acid/HATU/DiEA) and mixed for 1 min for pre-activation. The activated amino acid solution was added into the resin for coupling of 20 mins.

After coupling, the reaction solution was filtered, and the resin was washed twice with DMF, one time with DCM and one time with DMF. Negative amine test was observed for each coupling. The deprotection, coupling, washing and testing procedures were repeated for each cycle of amino acid attachment.

After synthesis, the peptides were subject to the final deprotection cycle and washed three times with DCM. The peptides were acetylated with acetic anhydride. In detail, 1:8:4 resin/acetic anhydride/DiEA were mixed and reacted for 45 mins. The acetylation was repeated once. The resin was then washed three times with DCM.

After acetylation, the peptides on the resin were mixed with the cleavage cocktail (90% TFA/2.5% Anisole/2.5% Milli Q H_2_O/2.5% TIPS/2.5% EDT by volume) and reacted for 3 hours. The cleaved peptide solution was drained into a clean round bottom flask. The TFA was blown off with nitrogen gas. Ice-cold diethyl ether was added into the peptide solution to triturate the peptide. The mixture was then centrifuged to obtain the white pellet of the crude peptide. The white pellet was washed once more with cold ether to remove the cleavage cocktail further. After that, the crude peptide was air-dried in the fume hood.

### Peptide Purification and Mass Spectrometry

The peptides were purified with the reverse phase high-performance liquid chromatography (HPLC) in a Waters instrument equipped with a C18 column (Waters). Milli-Q H_2_O with 0.05% w./v. TFA was mobile phase A and acetonitrile (Sigma Aldrich) with 0.05% w./v. TFA was mobile phase B. A general gradient of 5-50% B, 1.5%B/min, was used for peptide purification. The purified peptides were characterized of the molecular weight with a Bruker AutoFlex Speed MALDI ToF instrument or an Agilent qToF LC-MS instrument. Peptide purity post HPLC was checked with ultra performance liquid chromatography (UPLC, Waters).

### Peptide Sample Preparation for Folding

Purified peptide solutions were freeze-dried with a FreeZone 4.5L Freeze Dry System. Peptides solutions of 3 mM peptide in 20 mM MES buffer (Sigma Aldrich), pH 6.1±0.2 were made for self-assembly. The pH was adjusted with 1M NaOH and the pH value was confirmed with a pH meter (METTLER TOLEDO). The pH of the DES-1 peptide sample at pH 3.5 was adjusted with 1M HCl solution. Peptide solutions were annealed at 65 °C for 15 mins and then equilibrated at 4 °C for > 72 hrs before characterizations. The nanoribbon assembly from DES-1 peptide and the nanosheets structure from RD2 peptide require three weeks to form at room temperature.

### Circular Dichroism

The circular dichroism (CD) data were collected on a Jasco J-810 spectropolarimeter equipped with a Peltier temperature controller using a cuvette with 1 mm pathlength. Peptide solutions were diluted into 0.3 mM concentration with 20 mM MES buffer for data collection. The melting curves were collected from 5 °C to 65 °C with a heating rate of 10 °C/hour at 225 nm of each sample. The first-order derivatives of the melting curves were calculated with Savitzky-Golay smoothing algorithm, and temperature where the minimum derivative value appears was regarded as the melting temperature. The molar residue ellipticity (MRE) value was calculated with the equation, MRE = (θ × m)/(c × l × nr × 10) where θ represents the experimental ellipticity in millidegrees, m is the molecular weight of the peptide (g/mol), c is the peptide concentration (milligrams/milliliter), l is the path length of the cuvette (cm), and nr is the number of amino acid residues in the peptide.

### Size Exclusion Chromatography

The SEC data were collected in a Varian Star instrument equipped with a Superdex 200 Increase 10/300 GL column. The mobile phase was 100 mM MES buffer, and the flowrate was 0.75 ml/min. The peptide stock solution was diluted with 20 mM MES buffer and 100 uL of 0.6 mM peptide solution was injected for each measurement. The signal was monitored at 220 nm wavelength. The system was cooled and equilibrated at 10 °C for 45 mins before data collection to prevent denaturation of the supramolecular assemblies that are unstable at room temperature.

### Cryo-EM Characterization and Structure Reconstruction

Aliquots of the peptides were diluted with 20 mM MES buffer into 0.1 wt% for grid freezing. For sample dilution of DES-1 nanoribbon sample, the buffer was adjusted to be pH 3.5 before usage. All grids used for this study were on Lacey carbon 300 mesh copper grids. The grids were frozen with a Vitrobot™ Mark IV plunge freezer. Prior to the addition of the sample, the grids were cleaned using a Gloqube® Plus glow discharge system. Grids were imaged on a Glacios 200kV electron microscope with a cryo Autoloader, XFEG electron source and Falcon4 direct electron detector (1.2 Å/pixel). Processing of raw movies and subsequent image processing and reconstruction steps of images from cryo-EM were handled in cryoSPARC.^50^ Images were motion corrected and CTFs were estimated using the in house Patch Motion Correction and Patch CTF estimation software. For each dataset∼500 or so “particles” were selected using the manual particle picker function and these were then used to generate initial 2D class averages using the 2D classification program in cryoSPARC. For many samples multiple rounds of sorting based on 2D classification were performed to sort through obvious “junk” particles. For the SPA-human hexamers, DES-1 pentamers, and DES-1 hexamers ab initio reconstruction with multiple classes was performed with either C5 or C6 symmetry and the resulting “good” classes were then subjected to homogenous refinement with the same point group symmetry. For the DES-1 doublets the same basic protocol was followed except that C1 symmetry was imposed. Any density maps that were determined were then analyzed in UCSF Chimera^51^ and ChimeraX^52^.

### AFM Characterization

The atomic force microscope (AFM) characterization was performed in a Park NX20 AFM instrument using tapping mode, scanning rate 1.0 Hz. Measurements were performed using AFM tips with a force constant of 42 N/m and a resonance frequency of 320 kHz (NanoWorld). The peptide solution was diluted using 20 mM MES buffer to a final concentration of 0.3 mg/mL. For sample dilution of the DES-1 nanoribbon sample, the buffer was adjusted to be pH 3.5 before usage.10 uL of diluted peptide solution was drop cast to a freshly cleaved mica slide. After 5 mins of bonding, the solution was wicked away, and the mica slide was rinsed with 200 uL of fresh MilliQ H_2_O. The mica slide was air dry for 1h at room temperature before imaging.

### Dynamic Light Scattering

The samples were diluted ten folds with 20 mM MES buffer for measurement. The data were collected in a Malvern Zen 3600 Zetasizer instrument using Malvern ZEN0040 disposable cuvettes for size measurement. All measurements were done at 25 °C.

## Supporting information

Supplemental Experimental Data

## Notes

### Competing Interest Statement

The authors have declared no competing interest.

