## Supplemental Experimental Data for "Beyond the Triple Helix: Exploration of the Hierarchical Assembly Space of Collagen-like Peptides"

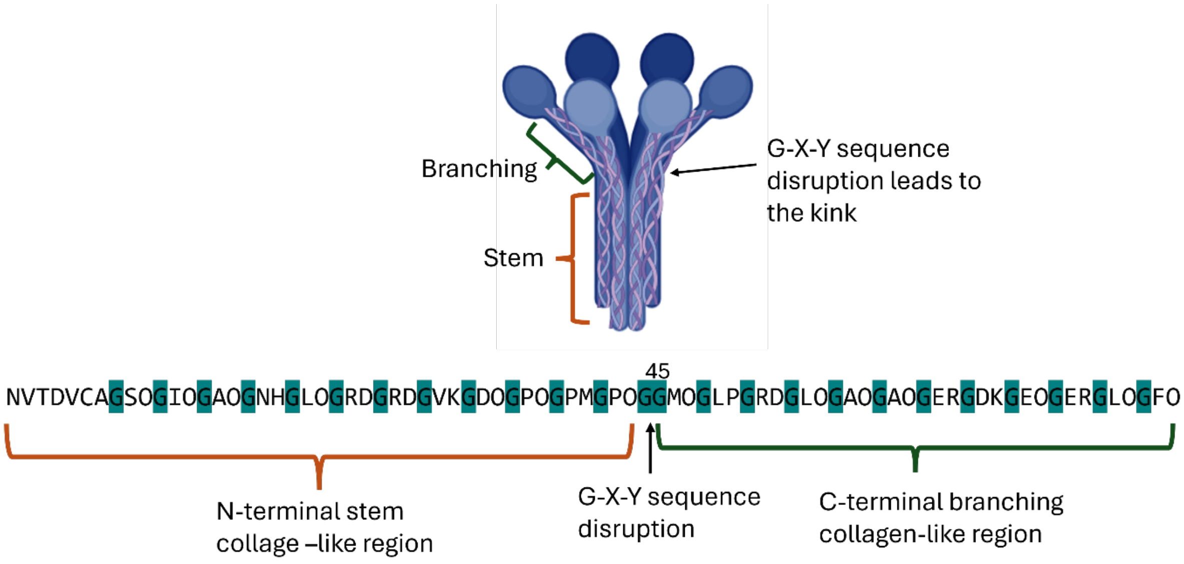


***Figure S1.*** *Illustration of amino acid sequence dissection of protein rat SP-A. The full-length protein sequence was obtained from the database.[1] "G" residue was highlighted. The amino acid sequence that conforms to the "G-X-Y" sequence pattern was considered as the collagen-like region. Disruption of the sequence pattern was observed at the 45$^{th}$ Gly residue (numbering starts after the signal peptide). This discontinuity was considered to cause the collagen triple helices to bend, forming the kink. Therefore, the N-terminal 1-44 residues were considered as the stem collagen-like region and the C-terminal 45-81 residues are considered as the branching collagen-like region.*

*Table S1 Peptide amino acid sequence and molecular weight information.*

| Peptides | Sequence | Expected $[M+H]^+$ | Observed $[M+H]^+$ |
|---|---|---|---|
| SPA-Rat | NVTDVAAGSOGIOGAOGNHGLOGRDGRDGVKGDOGPOGPMGPOG | 4206.5 | 4207.0 |
| SPA-Human | EVKDVAVGSOGIOGAOGNHGLOGRDGRDGVKGDOGPOGPMGPOG | 4282.0 | 4281.5 |
| SPA-bcb | AAGSOGIOGAOGNHGLOGRDGRDGVKGDOGPOGPMGPOG | 3479.7 | 3480.7 |
| SPA-R-P | NVTDVAAGSOGIOGAOGNHGLOGPDGRDGVKGDOGPOGPMGPOG | 4147.4 | 4147.9 |
| SPA-D-O | NVTDVAAGSOGIOGAOGNHGLOGROGRDGVKGDOGPOGPMGPOG | 4204.5 | 4204.4 |
| SPA-I-P | NVTDVAAGSOGPOGAOGNHGLOGRDGRDGVKGDOGPOGPMGPOG | 4191.4 | 4192.0 |
| SPA-I-S | NVTDVAAGSOGSOGAOGNHGLOGRDGRDGVKGDOGPOGPMGPOG | 4180.4 | 4180.9 |
| DES-1 | DAAGIOGPOGPOGLOGRDGRDGVOGPOGPOGPM | 3162.5 | 3163.0 |
| DES-H | DAAGIOGPOGPOGLOGPOGPOGVOGPOGPOGPM | 3063.3 | 3064.0 |
| RD1 | DAAGPOGPOGPOGRDGRDGRDGRDGPOGPOGPO | 3233.5 | 3233.7 |
| RD2 | DAAGPOGRDGRDGPOGPOGPOGPOGRDGRDGPO | 3233.5 | 3234.1 |
| RD2-no DAA | GPOGRDGRDGPOGPOGPOGPOGRDGRDGPO | 2976.0 | 2976.5 |

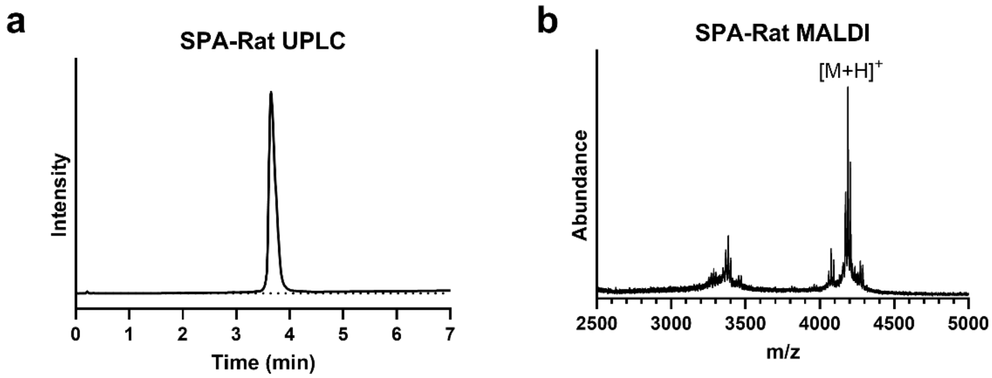


***Figure S2.*** *Peptide characterizations. a) Analytical HPLC (UPLC) of pure peptide SPA-Rat. The chromatographic method was run using a 5-35% B gradient with a 10% B/min ramping rate. b) MALDI mass spectrometry of peptide SPA-Rat.*

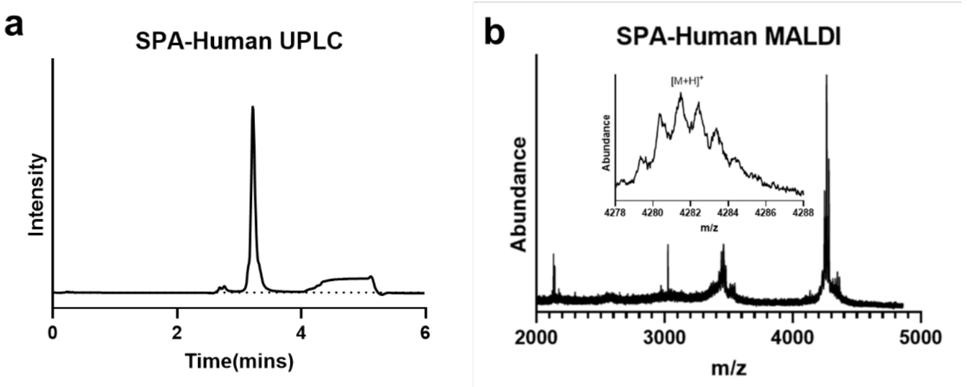


***Figure S3.*** *Peptide characterizations. a) Analytical HPLC (UPLC) of pure peptide SPA-Human. The chromatographic method was run using a 5-35% B gradient with a 10% B/min ramping rate. b) MALDI mass spectrometry of peptide SPA-human.*

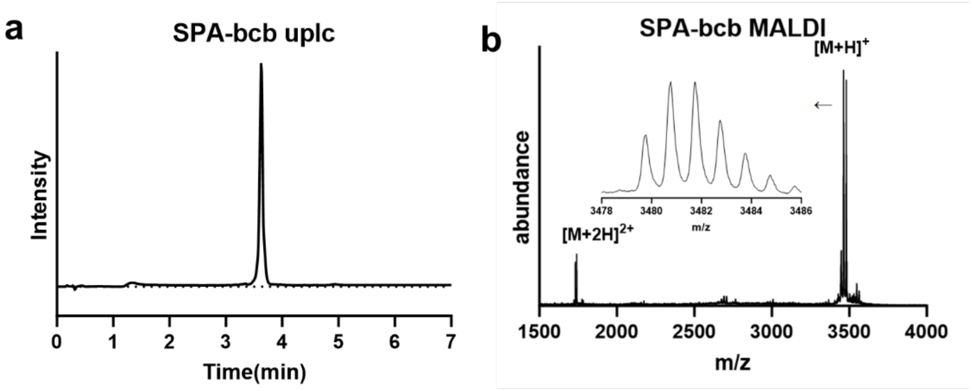


***Figure S4.*** *Peptide characterizations. a) Analytical HPLC (UPLC) of pure peptide SPA-bcb. The chromatographic method was run using a 5-35% B gradient with a 10% B/min ramping rate. b) MALDI mass spectrometry of peptide SPA-bcb.*

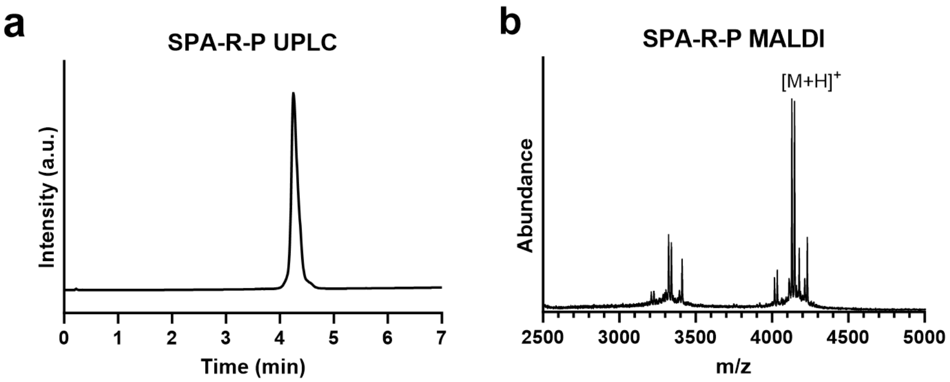


***Figure S5.*** *Peptide characterizations. a) Analytical HPLC (UPLC) of pure peptide SPA-R-P. The chromatographic method was run using a 5-35% B gradient with a 10% B/min ramping rate. b) MALDI mass spectrometry of peptide SPA-R-P.*

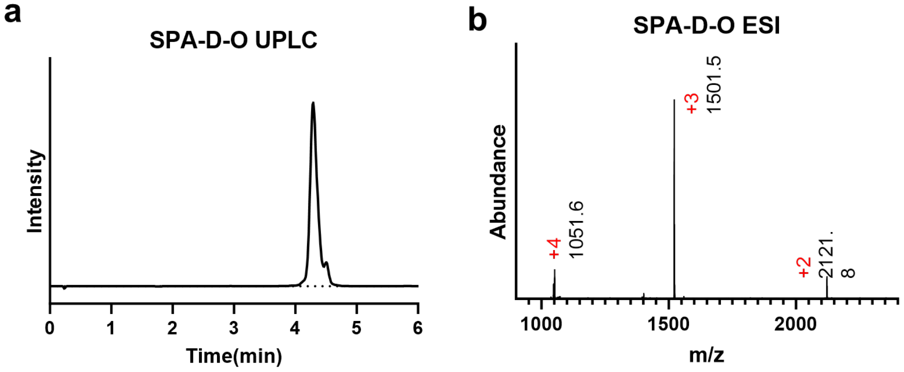


***Figure S6.*** *Peptide characterizations. a) Analytical HPLC (UPLC) of pure peptide SPA-D-O. The chromatographic method was run using a 5-35% B gradient with a 10% B/min ramping rate. b) LC-MS (ESI) mass spectrometry of peptide SPA-D-O.*

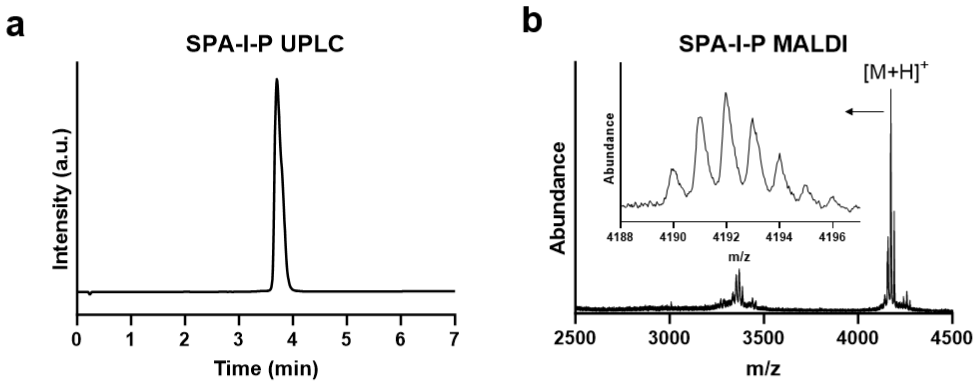


***Figure S7.*** *Peptide Characterizations. a) Analytical HPLC (UPLC) of pure peptide SPA-I-P. The chromatographic method was run using a 5-35% B gradient with a 10% B/min ramping rate. b) Mass spectrometry of peptide SPA-I-P.*

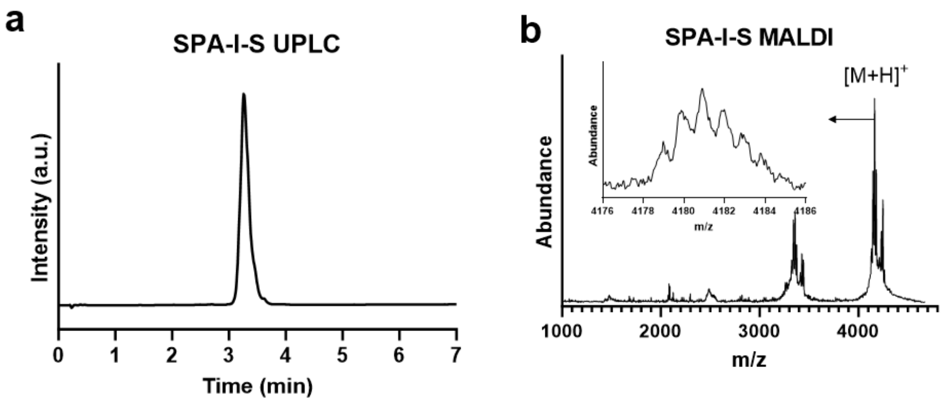


***Figure S8.*** *Peptide characterizations. a) Analytical HPLC (UPLC) of pure peptide SPA-I-S. The chromatographic method was run using a 5-35% B gradient with a 10% B/min ramping rate. b) Mass spectrometry of peptide SPA-I-S.*

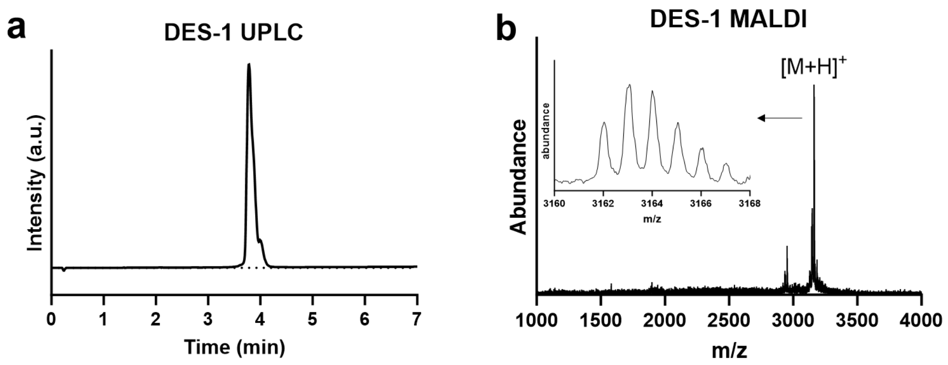


***Figure S9.*** *Peptide characterizations. a) Analytical HPLC (UPLC) of pure peptide DES-1. The chromatographic method was run using a 5-35% B gradient with a 10% B/min ramping rate. b) Mass spectrometry of peptide DES-1.*

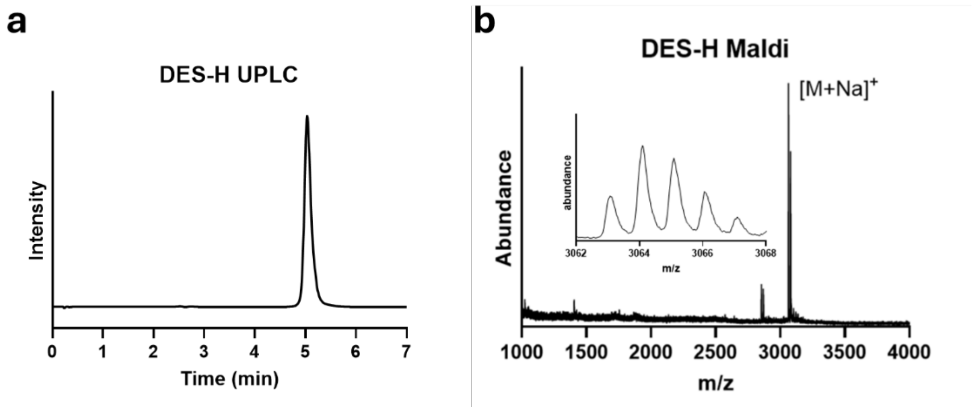


***Figure S10.*** *Peptide characterizations. a) Analytical HPLC (UPLC) of pure peptide DES-H. The chromatographic method was run using a 5-35% B gradient with a 10% B/min ramping rate. b) Mass spectrometry of peptide DES-H.*

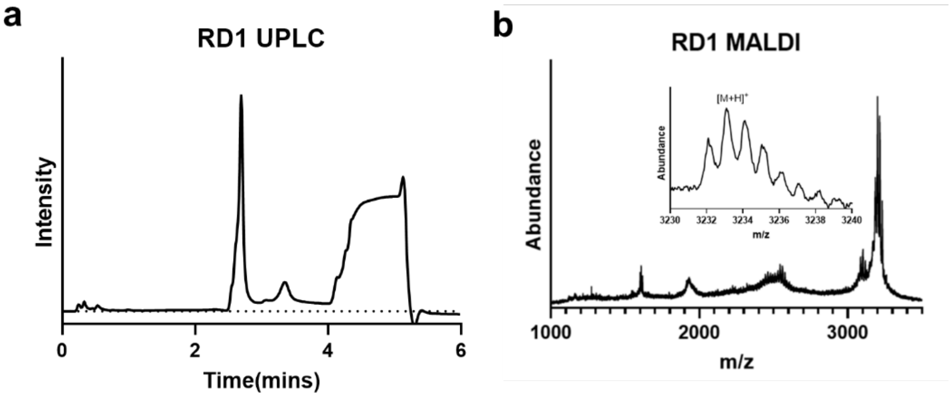


***Figure S11.*** *Peptide characterizations. a) Analytical HPLC (UPLC) of pure peptide RD1. The chromatographic method was run using a 5-35% B gradient with a 10% B/min ramping rate. The peak elutes after 4 min is in the wash peak and does not represent biomacromolecules. b) MALDI mass spectrometry of peptide RD1.*

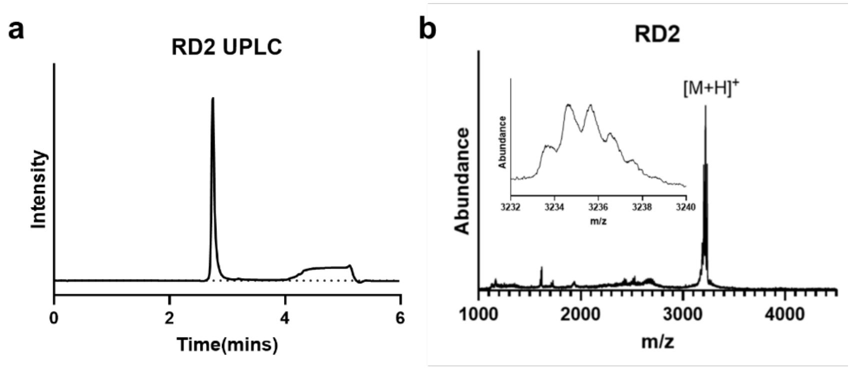


***Figure S12.*** *Peptide characterizations. a) Analytical HPLC (UPLC) of pure peptide RD2. The chromatographic method was run using a 5-35% B gradient with a 10% B/min ramping rate. b) MALDI mass spectrometry of peptide RD2.*

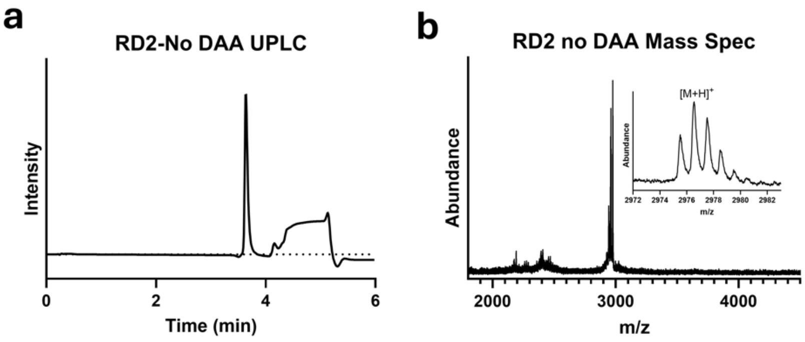


***Figure S13.*** *Peptide characterizations. a) Analytical HPLC (UPLC) of pure peptide RD2-No DAA. The chromatographic method was run using a 5-35% B gradient with a 10% B/min ramping rate. The peak elutes after 4 min is in the wash peak and does not represent biomacromolecules. b) MALDI mass spectrometry of peptide RD2-No DAA.*

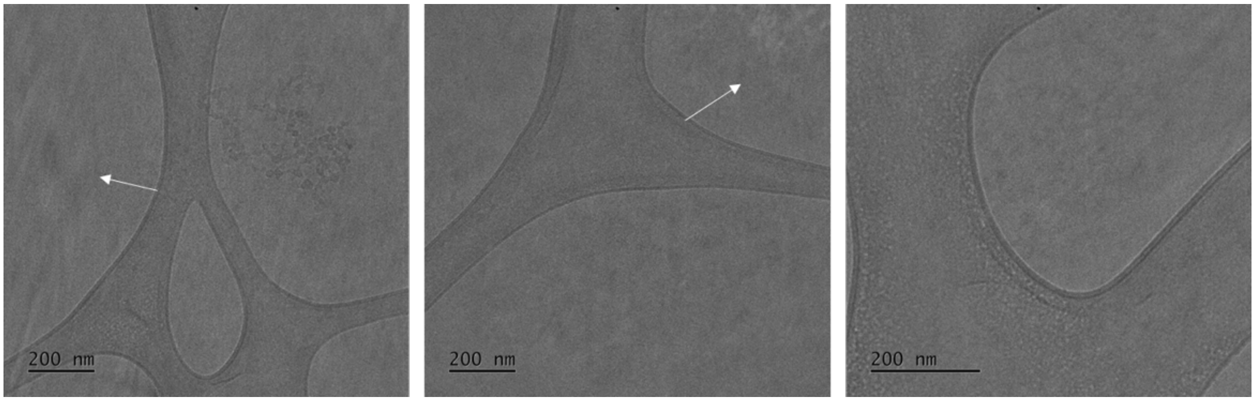


***Figure S14.*** *Cryo-TEM characterization of SPA-bcb aggregates. White arrows highlight the amorphous features of the aggregates. No uniform assembly was observed.*

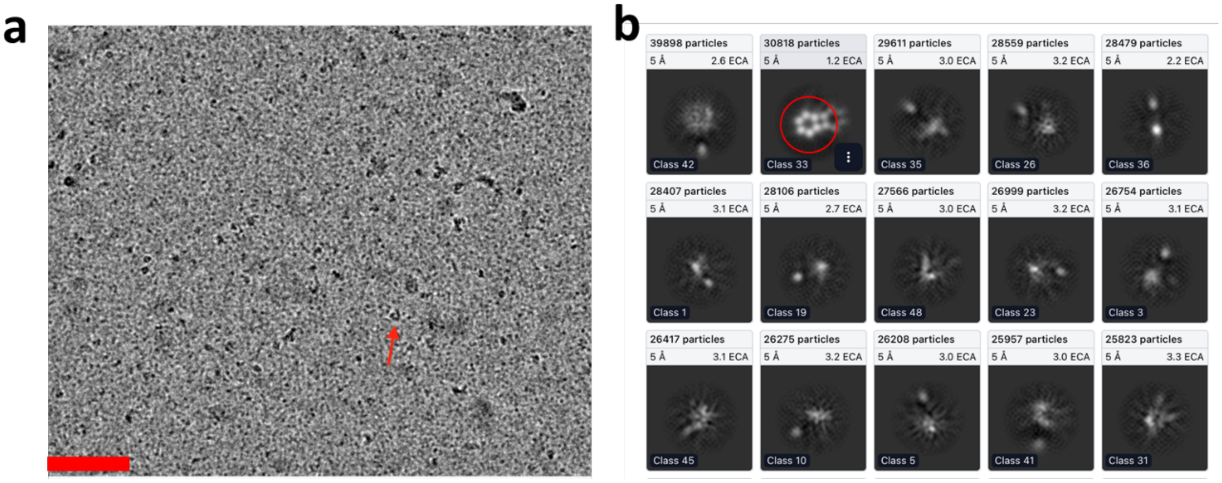


***Figure S15.*** *a) representative cryo-EM image of SPA-Rat peptide. b) 2D class averaging of the cry-EM images collected on peptide SPA-Rat.*

***Table S2** Parameters of the models constructed from the cryo-EM data.*

| Reconstruction | EMDB number | Resolution (Å) |
|---|---|---|
| DES-1 Doublet | EMD-44405 | 10.2 |
| DES-1 Octadecamer (C6) | EMD-44406 | 5.3 |
| DES-1 Pentadecamer (C5) | EMD-44409 | 5.0 |
| SPA-human octadecamer | EMD-44418 | 5.4 |

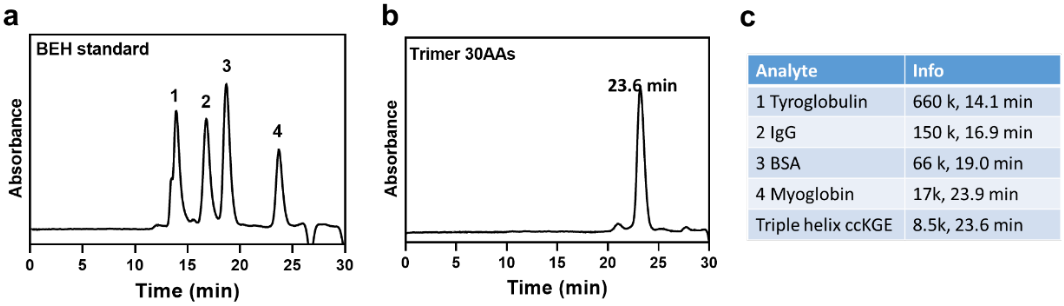


***Figure S16.*** *SEC characterization of protein standards in the Superdex 200 Increase column with a flow rate of 0.75 mL/min. a) SEC of the commercial BEH standard from Waters. B) SEC of a covalently captured collagen trimer with a molecular weight of approximately 8.5 kDa. The trimer was made in our previous work.[2] c) molecular weight information of the standard protein species.*

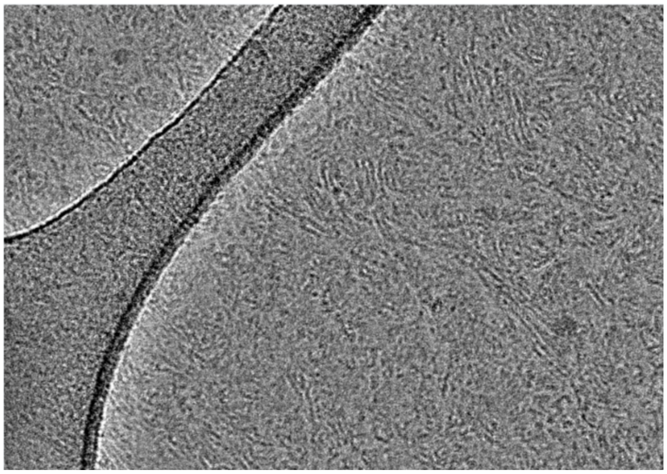

***Figure S17.*** *Additional representative cryo-EM image of SPA-Human assembly.*

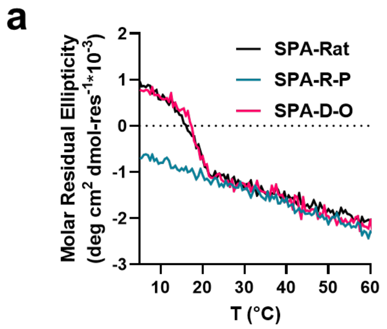


***Figure S18.*** *CD melting results of SPA peptides. a) CD melting curves of SPA-Rat, SPA-R-P and SPA-D-O peptide mutants. b) CD melting curves of SPA-Rat, SPA-I-P, and SPA-I-S samples.*

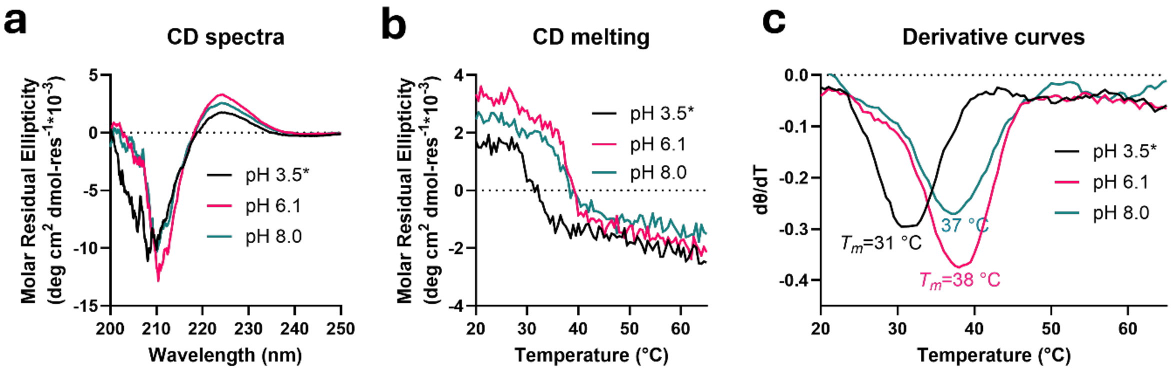


***Figure S19.*** *CD characterization of DES-1 peptide folded at pH 3.5, 6.1 and 8.0. After 3 weeks of equilibration, the peptide folded in pH 3.5 precipitated and the solution became turbid which may impact the quality of CD measurement. a) CD spectra. b) CD melting curves. Signal was monitored at 224 nm. c) The first-order derivative curves of the melting curves in b).*

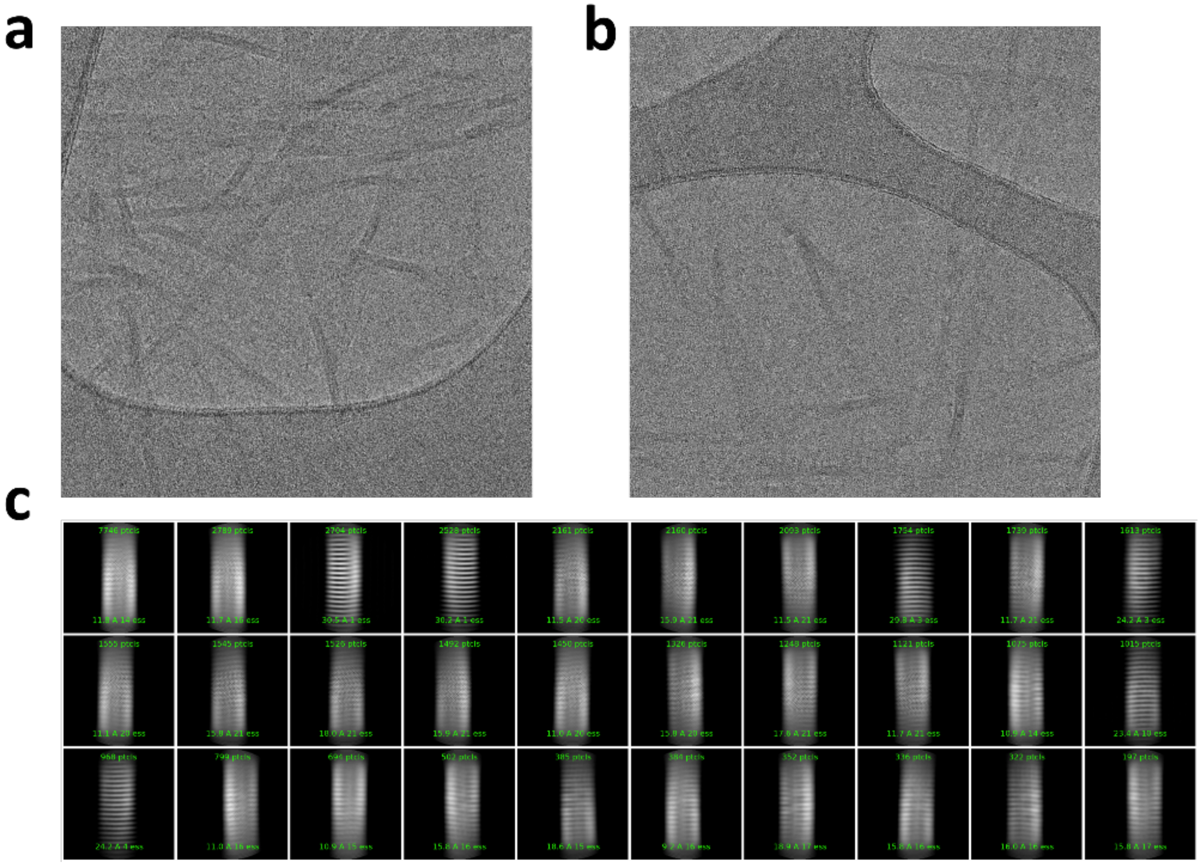

***Figure S20.*** *Additional representative cryo-EM results of DES-1 peptide at pH 3.5.*

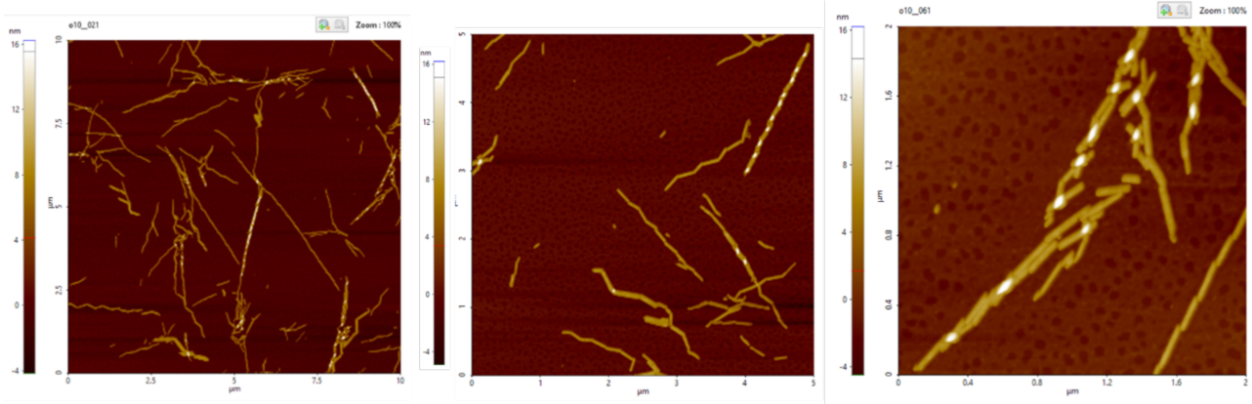

***Figure S21.*** *Additional AFM results of DES-1 peptide assembled at pH 3.5.*

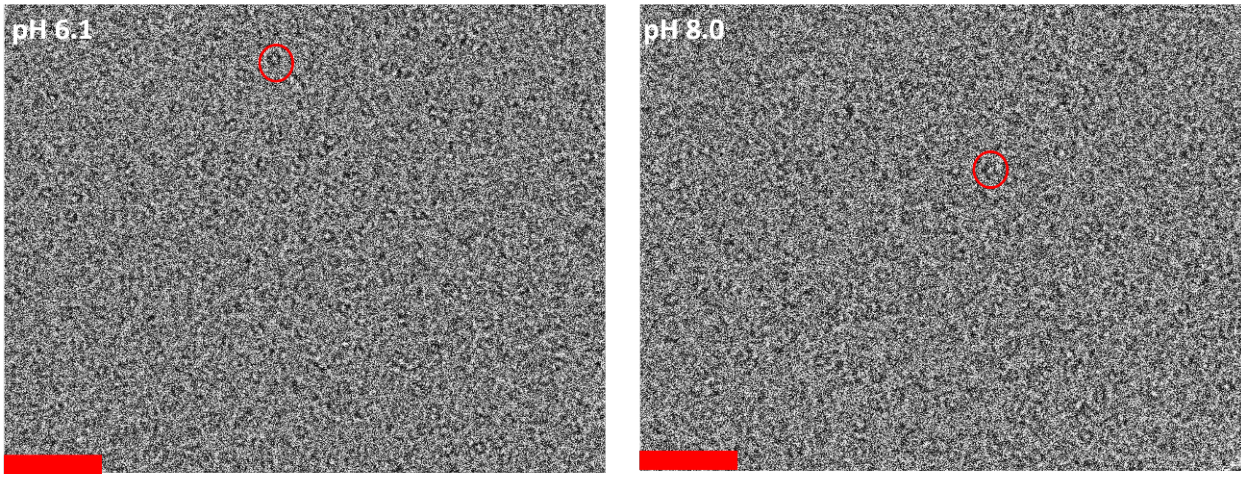


***Figure S22.*** *Representative raw cryo-EM image of DES-1 peptide at pH 6.1 and 8.0. The scale bars are 50 nm.*

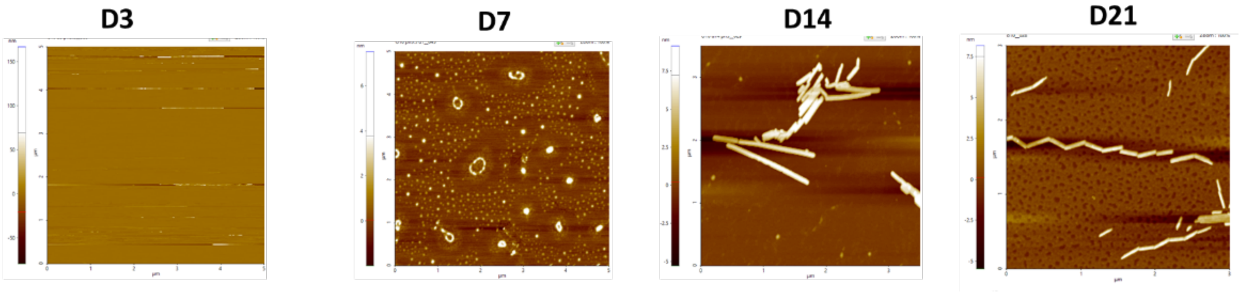


***Figure S23.*** *AFM results of DES-1 peptide assembled at pH 3.5 after different time of equilibration.*

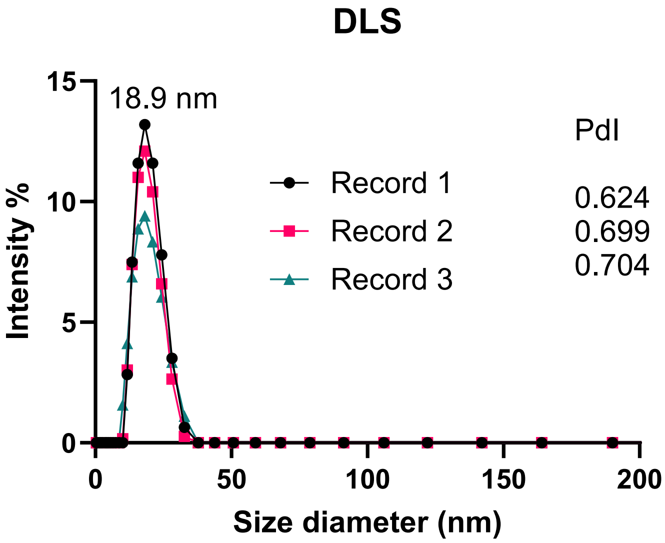


***Figure S24.*** *Dynamic light scattering of RD1 nanoparticles.*

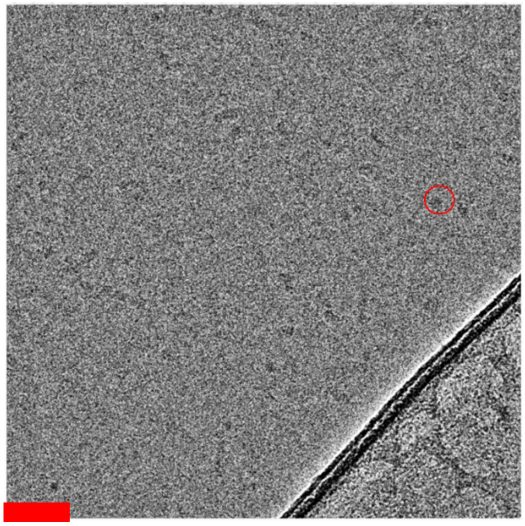

***Figure S25.*** *Representative cryo-EM image of RD1 peptide assembly. The red circle highlights one of the RD1 protein nanoparticles. Scale bar is 50 nm.*

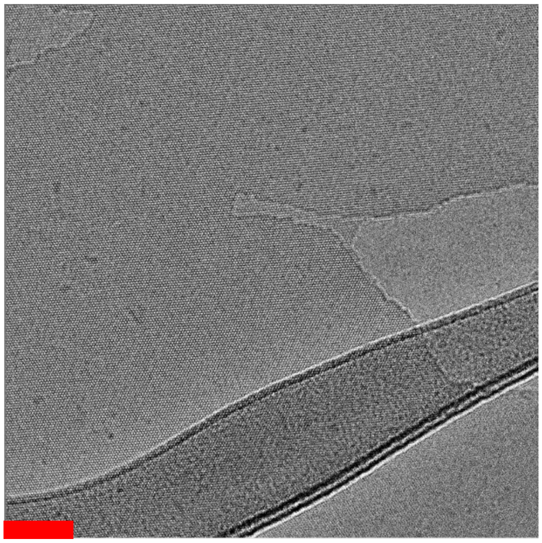

***Figure S26.*** *Additional representative cryo-EM image of the RD2 nanosheets. The scale bar is 50 nm.*

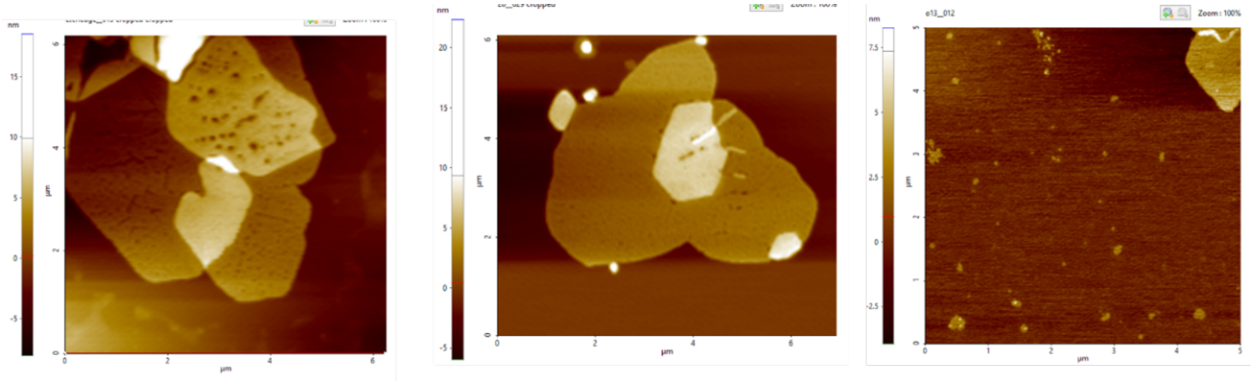


***Figure S27.*** *Additional AFM characterization results of RD2 two-dimensional nanosheets.*

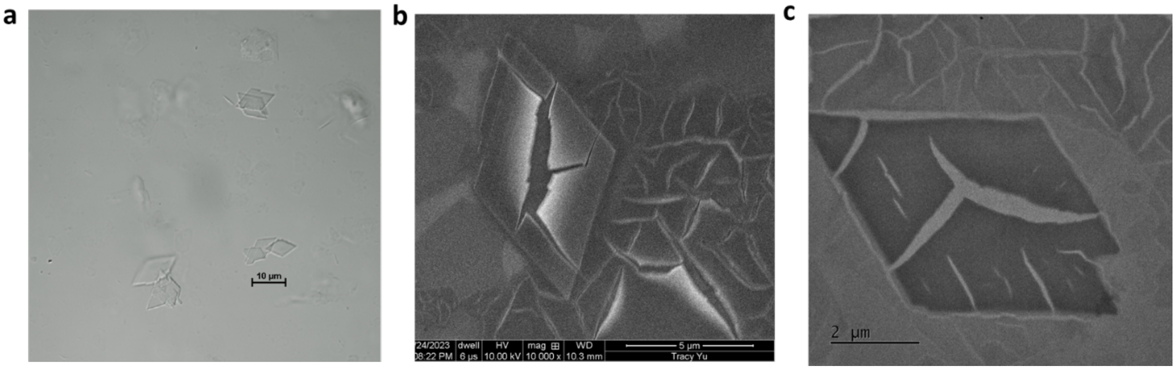


***Figure S28.*** *a) optical microscope of RD2-No DAA peptide. Microcrystals were observed. b) SEM of RD2-No DAA. c) TEM of RD2-No DAA microcrystals.*

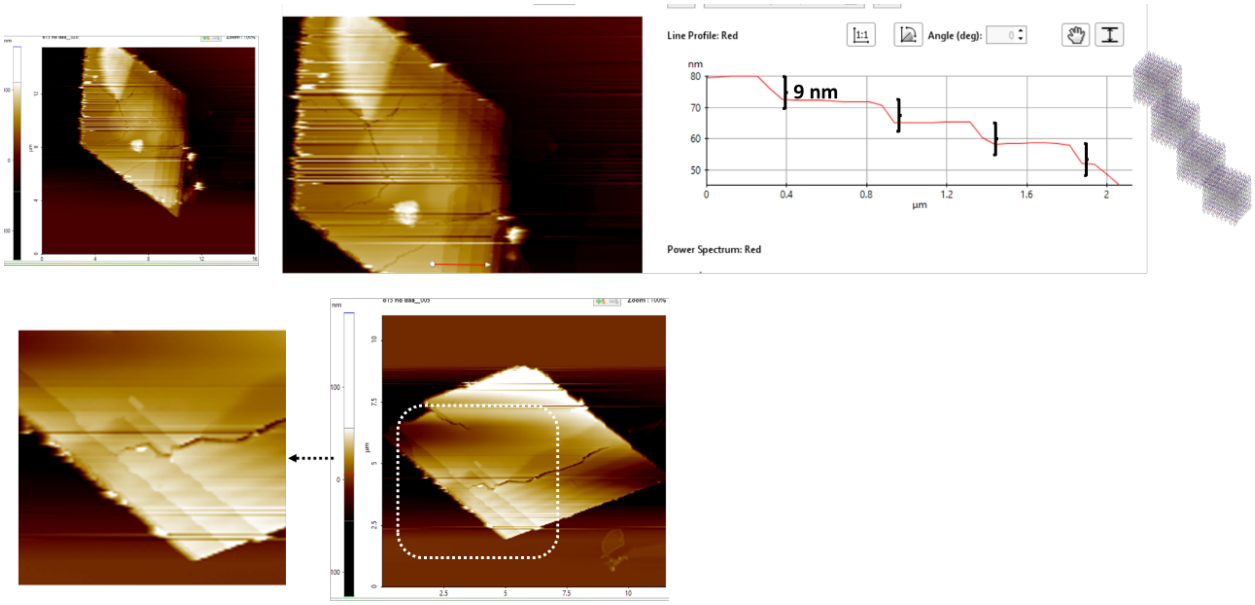


***Figure S29.*** *AFM characterization of RD2-No DAA microcrystals.*
